# The importance of the photosynthetic Gibbs effect in the elucidation of the Calvin-Benson-Bassham Cycle

**DOI:** 10.1101/200105

**Authors:** Oliver Ebenhöh, Stephanie Spelberg

## Abstract

The photosynthetic carbon reduction cycle, or Calvin-Benson-Bassham Cycle, is now contained in every standard biochemistry textbook. Although the cycle was already proposed in 1954, it is still subject of intense research, and even the structure of the cycle, i.e. the exact series of reactions, is still under debate. The controversy about the cycle’s structure was fuelled by the findings of Gibbs and Kandler in 1956 and 1957, when they observed that radioactive ^14^CO_2_ was dynamically incorporated in hexoses in a very atypical and asymmetrical way, a phenomenon later termed the ‘photosynthetic Gibbs effect’. Now, it is widely accepted that the photosynthetic Gibbs effect is not in contradiction to the reaction scheme proposed by Calvin, Benson and Bassham, but the arguments given have been largely qualitative and hand-waving. To fully appreciate the controversy and to understand the difficulties in interpreting the Gibbs effect, it is illustrative to illuminate the history of the discovery of the Calvin-Benson-Bassham Cycle. We here give an account of central scientific advances and discoveries, which were essential prerequisites for the elucidation of the cycle. Placing the historic discoveries in the context of the modern textbook pathway scheme illustrates the complexity of the cycle and demonstrates why especially dynamic labelling experiments are far from easy to interpret. We conclude by arguing that only mathematical models based on a sound theory are capable of resolving conflicting interpretations and providing a consistent quantitative explanation.

## Introduction

The Calvin-Benson-Bassham (CBB) Cycle (1,2) – often simply termed ‘Calvin Cycle’ or ‘Calvin-Benson Cycle’ (3) or, as preferred by James Alan Bassham, the ‘Photosynthetic Carbon Reduction (PCR) Cycle’ (4) – is undoubtedly one of the most important biochemical pathways on earth. Plants and many other photosynthetic organisms employ it to fix carbon dioxide and reduce it to sugars. The key enzyme RuBisCO, which catalyses the carboxylation reaction, is probably the most abundant enzyme on earth (5,6), responsible for over 99% of global carbon dioxide fixation (7). Not surprisingly, the biochemical mechanisms of inorganic carbon fixation have been an intense subject of scientific investigations since the early days of metabolic research.

Since first suggested by the German chemist Adolf Baeyer in 1870 (8), it was widely assumed that the primary product of carbon fixation should be formaldehyde, formed by photo-excited chlorophylls, and that formaldehyde molecules undergo subsequent polymerisation to sugars (9). This hypothesis even received experimental support by Klein and Werner in 1926 (10) using the chemical dimedon (dimethylhydroresorcinol), which allows detecting even small quantities of formaldehyde. However, when repeating these experiments to address a few unsolved questions, such as the observation that formaldehyde was mainly found in the extracellular medium, Barton-Wright and Pratt could show in 1930 (11) that formaldehyde production was an experimental artifact and ruled out that it is connected to photosynthesis. Lacking a plausible alternative theory, the formaldehyde hypothesis was nevertheless still widely accepted.

### The interdisciplinary breakthrough: radioactive isotopes

It took the intuition of interdisciplinary scientists to apply techniques from nuclear physics to photosynthesis research to falsify the formaldehyde hypothesis. With the invention and actual construction of the first cyclotron (12) at the University of California, Berkeley, in 1932, it became for the first time possible to generate high-energy particles and use them to produce radionuclides. Applying this new technology to produce the radioactive isotope carbon-11 (a positron emitter with a half-life of 20 minutes) opened dramatically new perspectives, because it allowed marking single carbon atoms and follow their fate, regardless in which molecules these atoms are bound and which chemical or biochemical reactions they undergo. The scientists have developed "an eye, which could look into the plant cells" (13). Using carbon-11, Samuel G. Ruben and Martin Kamen could show that the primary product of carboxylation must contain a carboxyl group (14). These experiments have not only started the long and tedious elucidation of the complete CBB cycle, but they also represent the birth of using radiolabelling techniques in biological research. Owed to the short half-life of carbon-11, which limited the possible experiment time to just a few hours, it was not suited to reveal more details about the series of reactions now known as the CBB cycle. Experimenting with different targets bombarded with high-energy particles in the cyclotron, Ruben and Kamen discovered different ways to produce carbon-14 as early as 1941 (15). Partly due to the low beta emission and low specific activity, but probably mainly due to the obvious hindrances to carry out research during World War II, they never used it in photosynthetic experiments (16). However, with the discovery of the long-lived isotope carbon-14 (half-life of ~5700 years) they set the foundation for the major breakthrough of Melvin Calvin and his team after the end of the war.

### Melvin Calvin’s quest for the photosynthetic carbon fixation pathway

Only after World War II, starting in 1945, photosynthesis experiments with carbon-14 were systematically carried out by Melvin Calvin and Andrew A. Benson, who operated their photosynthesis research laboratory adjacent to the cyclotron, in which carbon-14 could be produced (16,17). Benson first confirmed with carbon-14 what Ruben had conjectured already with carbon-11, that the primary product of carboxylation contains a carboxyl group. Experiments with preilluminated *Chlorella* and *Scenedesmus* algae provided evidence that the first product of carbon fixation is phosphoglyceric acid (PGA) (13,18). Refining the technique of two-dimensional paper chromatography (13), and varying the illumination time, during which *Chlorella* and *Scenedesmus* were fed with radio-labelled carbon dioxide, it became possible to identify the organic compounds containing the radioactive carbon atoms.

Nevertheless, the long-standing question of the nature of the acceptor of the carbon atom from carbon dioxide was still unsolved. Observations by Calvin that the pool sizes of ribulose 1,5-bisphosphate (RuBP) and phosphoglyceric acid changed in a reciprocal fashion when algae were subjected to changes in light intensity (19) gave strong hints that ribulose 1,5-bisphosphate was indeed the substrate providing the 2-carbon backbone to which the carbon atom from carbon dioxide was fixed. In their seminal paper from 1954, Melvin Calvin and his co-workers proposed their famous scheme, which is now known as the Calvin-Benson-Bassham Cycle. There they confirmed the reciprocal behavior of ribulose 1,5-bisphosphate and phosphoglyceric acid also under changes in carbon dioxide concentration. Apparently, they had no doubt regarding the nature of the carbon acceptor and it is therefore even more remarkable that the enzyme catalysing the formation of PGA from RuBP and carbon dioxide was isolated and characterised (20) only two years later by Weissbach, Horecker and Hurwitz. Interestingly, it took another year until it was realised by Dorner and Kahn (21) that this enzyme, termed ‘carboxylation enzyme’ by Horecker and ‘carboxydismutase’ by Calvin, was actually the same as the ‘Fraction 1 protein’, already isolated in 1947 at CalTech (Pasadena) by Wildman and Bonner by ammonium sulfate fractionation (22). The now commonly used name RuBisCO was only coined more than 20 years later by David Eisenberg in a talk at a symposium honouring Wildman (23).

However, even the strong indication Calvin and coworkers had regarding the nature of the carbon dioxide acceptor molecule, the problem was still open by which series of reactions this substrate could be recycled. Here, the pioneering biochemical research by Racker and Horecker set the foundation to derive a plausible reaction scheme. Both researchers investigated the series of reactions, now known as the Pentose Phosphate Pathway, by which pentose phosphates are converted into hexose phosphates. In the early 1950s, both groups independently published the action of the enzyme transketolase (a name coined by Racker), isolated from rat liver (24), yeast (25) and spinach (26), demonstrating that it catalyses the transfer of two carbons from a keto-sugar to an aldo-sugar. Simultaneously, Horecker could isolate and characterise the enzyme aldolase from rat liver (27), which together with his isolation of transaldolase from rat liver and yeast (28) forms the core of the Pentose Phosphate Pathway. Knowing of these results, and excluding a number of chemically feasible alternatives by a serious of careful experiments with short pulses of radioactive carbon dioxide, Melvin Calvin and coworkers finally proposed their famous reaction scheme in 1954 (2). However, in contrast to the carboxylation, the authors were apparently uncertain regarding the existence and activity of transketolase in chloroplasts, as is nicely illustrated by the question marks at the respective locations in the proposed scheme in Fig. 7 of the 1954 publication.

### Challenging Calvin’s scheme: The photosynthetic Gibbs effect

The 1954 paper by Melvin Calvin and his collaborators is an illustrative example of rigorous scientific work. Experimental evidence is provided in various forms, foremost in short-pulse ^14^CO_2_ labelling experiments to identify in which sugars the radioactive label is found, and - by chemical degradation – also the position of the labels in pentoses and heptoses was determined. By logic reasoning various potential pathways were excluded, so that the paper concludes with the scheme of the Calvin-Benson-Bassham Cycle as a logically derived, plausible hypothesis. However, the authors are also critical with their own results and suggest further experiments how the reaction scheme could be verified. Here is where Martin Gibbs enters the scene. Appointed in 1947 as junior scientist to the Brookhaven National Laboratory, his task was to synthesise radiocarbon labelled sugars and to provide them to other researchers. Because the only available compound with radioactive carbon at that time was barium carbonate, Martin Gibbs (chemist by training, PhD in botany) thought it natural to produce these sugars from radioactive carbon dioxide using photosynthesis. One of the early problems he was facing was to localise the radioactive carbon in the produced sugars. Together with Irwin C. Gunsalus, this problem was solved exploiting the unusual fermentation route of *Leuconostoc mesenteroides*, which degrades glucose into carbon dioxide (from carbon 1), ethanol (carbons 2 and 3) and lactic acid (carbons 4, 5 and 6) (29). This fermentation pathway allowed a unique determination of the positions of the radioactive carbon atoms, a decisive advantage over the degradation by *Lactobacillus casei*, introduced by Wood, Lifson and Lorber in 1945 (30), which only allowed a separation into 2-carbon groups (3-4, 2-5, 1-6). The *Leuconostoc* method soon became a standard for many labs interested in carbon metabolism. Bernard Horecker initially used this method to investigate the conversion of pentose phosphates into hexose phosphates in rat liver (31) by applying pentose phosphate that was labelled either in position 1 or in positions 2 and 3. The results gave rise to suggesting a series of reactions, now known as the Pentose Phosphate Pathway, in which reactions catalysed by transketolase, transaldolase and aldolase convert 6 pentoses in 5 hexoses. In the same year, Gibbs and Horecker studied the same pathway in extract from pea roots and leaves (32). While they found that the root extract showed essentially the same isotope label pattern as rat liver extract, the experiments with leaf extract were significantly different. Aware of the ongoing studies by Calvin to unravel the path of carbon in photosynthesis, who have demonstrated that ribulose, sedoheptulose, fructose and glucose phosphates are rapidly labelled during photosynthesis (33), he concluded that the path to recycle pentoses into hexoses is of significant importance in photosynthesis.

Otto Kandler, a German botanist from Munich, was sceptical about the carbon fixation scheme proposed by Calvin and visited Martin Gibbs to test the proposed pathway with the newly developed *Leuconostoc* method. Before establishment of this new method, various researchers (34–36) reported that photosynthetically derived hexoses are rapidly labelled in the 3,4 positions, while the label appeared later in the 2,5 and 1,6 positions, where sometimes an equal and sometimes an unequal label intensity was observed (37). Comparing these results to experiments of carbon fixation in the dark (37) or non-photosynthetic tissues like rat liver (30), in which the label was almost exclusively found in the 3,4 positions led to the assumption that the initial steps of carbon fixation are identical in the light and dark, and that the presence of light was responsible only for the further distribution of the labels; a hypothesis that was rapidly refuted after the initial experiments of Kandler and Gibbs.

Applying the new *Leuconostoc* method, Kandler and Gibbs observed, to their great surprise, that the distribution of labels in glucose phosphate (38) and starch derived glucose (39) formed during photosynthesis was in fact very asymmetric. Regardless of the precise experimental conditions and the botanical origin of the hexoses or hexose phosphates investigated, they found a persistent asymmetry in that the 4-carbon was always labelled before the 3-carbon, while the 1-carbon was labelled stronger than the 6-carbon and the 2-carbon stronger than the 5-carbon. This surprising finding of asymmetric label incorporation during photosynthetic carbon fixation is now often referred to as the ‘Gibbs effect’, which, as stressed by Chrispeels in his biographical memoirs of Martin Gibbs, “is yet to be explained” (40).

Kandler and Gibbs observed these asymmetries in various algae and plant species, but the exact order, in which carbons 1, 2, 5 and 6 are labelled, differed. So how can the common features, such as the 4/3, 1/6 and 2/5 asymmetries, be interpreted and how can it be understood, which processes determine the exact labelling dynamics?

Some aspects can indeed be simply explained. Bassham noted already in 1964 (41) that the asymmetry in the 3 and 4 position can be explained by different pool sizes of the triose phosphates. Radioactive carbon from carbon dioxide is incorporated into glyceraldehyde phosphate (GAP) in the carboxyl carbon (carbon 1). GAP is isomerised to dehydroxy acetone phosphate (DHAP) in a reversible reaction, which is close to equilibrium (42). However, the DHAP pool is approximately 20 times larger than the GAP pool. Therefore, the fraction of radioactive carbon atoms in position 3 of DHAP (which results from the 1-carbon of GAP after isomerisation) increases considerably slower than the label in GAP. During condensation of GAP and DHAP by aldolase, the 1-carbon of GAP becomes the 4-carbon of fructose 1,6-bisphosphate (FBP), while the 3-carbon of DHAP enters position 3. Since during the further conversion to hexose phosphates and hexoses stored in starch, the position of carbon atoms is unchanged, it is obvious that Gibbs and Kandler always observed a higher radioactivity in position 4 compared to position 3.

In contrast, explaining the 1/6 and 2/5 asymmetries is far more challenging. Although Bassham explained the observed pattern by the ‘reversibility of transketolase’ (41), it remains difficult to find verbal explanations for the exact appearance of labels in the various intermediates of the Calvin-Benson-Bassham Cycle. Indeed, Trebst and Fiedler (43) argued in 1962 that the observations by Kandler and Gibbs ‘are in full agreement’ with the proposed reaction scheme, whereas Stiller concludes in the same year (44) that the ‘entire pathway from triose phosphate to pentose phosphate … is inadequate in a wide selection of organisms’.

### The rediscovery of quantitative biology as an interdisciplinary science

Traditionally, biological researchers have often sought theoretical explanations of their observations, and mathematical formulae have been commonly found in many publications to support interpretations. Also Bassham has employed quantitative arguments in his 1964 publication (41) in an attempt to explain the asymmetric label distribution observed by Gibbs and Kandler (38,39). His mathematical considerations are in fact simplifying mathematical models, which attempt to explain observations based on theoretical considerations. Quantitative predictions can best be generated by mathematical models, which are based on a sound theory. At the Metabolic Pathway Analysis conference in Bozeman, MT in 2017, we have presented preliminary results of a dynamic model that simulates the label incorporation dynamics of the CBB cycle. The model incorporates all reactions originally proposed by Calvin and coworkers (2), and the parameters have been chosen, such that the model exhibits experimentally determined steady-state pool sizes (42). Together with the thermodynamic properties of the reactions, the pool sizes determine the degree of reversibility, in particular for those reactions catalysed by transketolase, aldolase, isomerases and epimerases. Simulating the label distribution involved definition of dynamic variables representing all possible labelling patterns for all intermediates. For example, 2^3^=8 variables describe the patterns of GAP, while 2^7^=128 versions of sedoheptulose 1,7-bisphosphate (SBP) are included. Reactions were included reflecting the exact carbon transition maps for all transformations, resulting in 3904 isotope-specific rate equations. Under the chosen conditions, the model predicts the dynamic appearance of radioactive labels in hexose phosphate as shown in Fig. 1. The experimentally observed orders (4 before 3, 1 before 6, and 2 before 5) are clearly reproduced. While it should be stressed that these results are only reflecting one particular experimental condition, and generality still remains to be demonstrated, the theoretical results indicate that the consistently observed asymmetries are a direct consequence of the structure and the thermodynamics of the CBB cycle.

**Figure 1:**
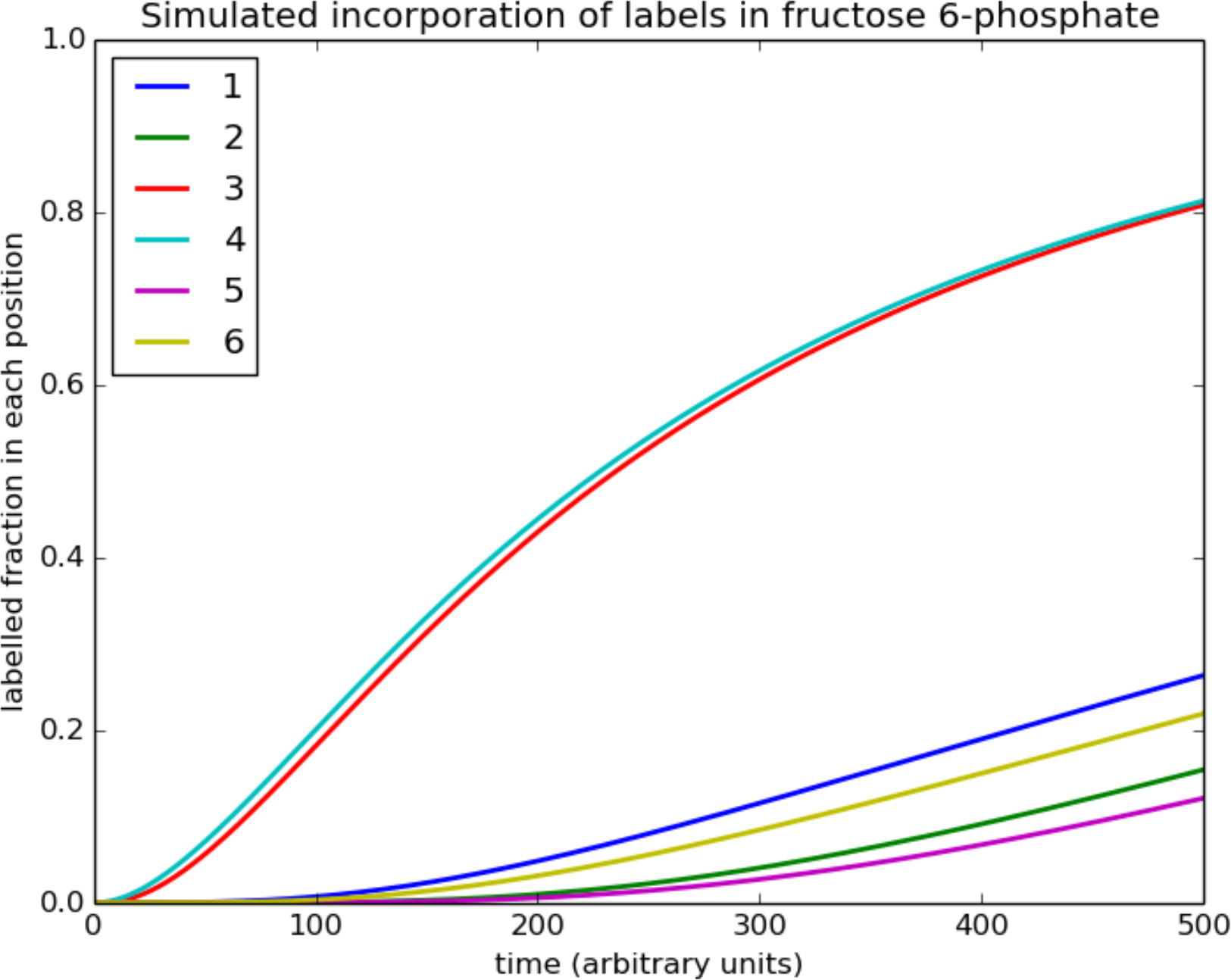
Simulated dynamic incorporation of radioactive carbon during photosynthesis. Displayed is a simulated time course of the incorporation of radioactive carbon dioxide into the CBB cycle intermediate fructose 6-phosphate. As observed in the Gibbs effect, labels are introduced in position 4 before position 3, and appear earlier in position 1 (2) than in 6 (5).

So is the scheme of Calvin finally proven? Not by a long shot!

One of the key enzymes in the recycling process from triose phosphates to pentose phosphates is transketolase, which – as already subsumed by Calvin in 1954 – catalyses two key steps: the transformation of a triose phosphate and a hexose phosphate into a tetrose phosphate and a pentose phosphate, and of a triose phosphate and a heptose phosphate into two pentose phosphates. Both reactions occur according to the same scheme: carbons 1 and 2 from a ketose are transferred to an aldose. Considering that the catalysed reactions are readily reversible (see e.g. (42)), the donor ketose is a pentose, hexose or heptose, while the acceptor aldose is a triose, tetrose or pentose. There is no convincing argument why the four (two reactions in two directions) transformations should be the only ones transketolase can catalyse. Why should not all nine possible combinations of ketose and aldose phosphates serve as substrates? In fact, using radiocarbon labelling, Clark showed *in vitro* that rat liver transketolase catalyses also the neutral exchange reaction erythrose 4-phosphate (E4P) +glucose 6-phosphate (G6P) <=> G6P+E4P (45), indicating clearly that the specificity of transketolase is not limited to the two reactions in the classical CBB cycle scheme. Moreover, it has long been debated whether longer sugars, such as 8 carbon octoses, play a role in the recycling reactions. In 1978 Williams proposed an alternative reaction scheme for the non-oxidative pentose phosphate pathway in rat liver involving five additional intermediates (arabinose 5-phosphate, <u>D-glycero-D-ido-octulose1,8-bisphosphate</u>, <u>D-glycero-D-altro-octulose1,8-bisphosphate</u>, altro-heptulose1,7-bisphosphate and manno-heptulose 7-phosphate) (46). In 2006, Williams and Flanigan demonstrated transketolase activity for C8 sugars and measured the distribution of labelled ^13^CO_2_ in these octoses using a new Gas Chromatography and Electron Ionization Mass Spectrometry methodology combined with Selected Ion Monitoring (GC/EIMS/SIM) (47,48). Therefore, transketolase presents a far more complex picture than originally assumed by Calvin. Seemingly complex, all possible activities by transketolase can easily be explained by the basic mechanism of transferring a 2-carbon group from a ketose to an aldose phosphate. All possible combinations can be compactly depicted by the scheme in Figure 2. Here, columns represent the accepting aldo-sugar phosphates (lengths 3 to 6) and the rows represent donating keto-sugar phosphates (lengths 5 to 8). Every field represents a possible substrate pair, and the resulting products are found in the field diagonally opposite. Considering sugars with up to 8 carbons results in a total of 16 possible reactions, four of which are neutral. Potentially, all these reactions will contribute to distributing the labelled around the different positions in the intermediate sugar phosphates.

**Figure 2A:**
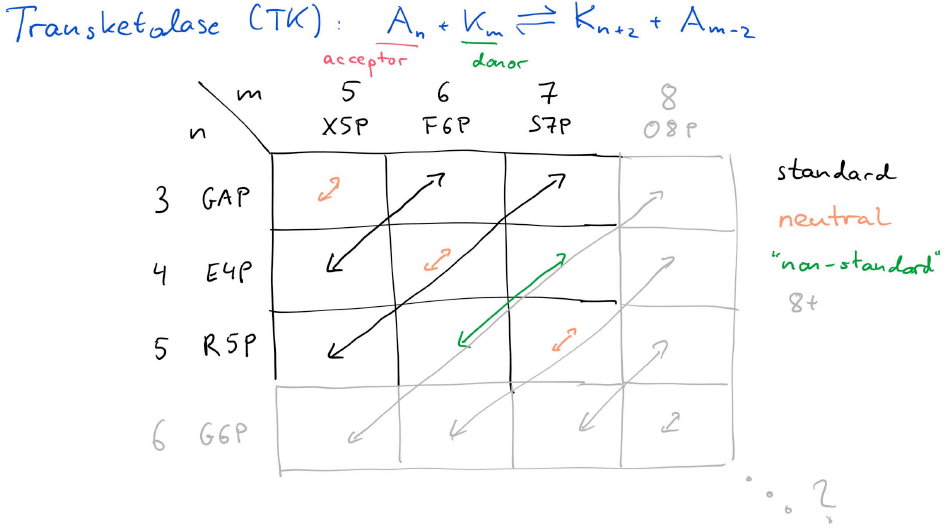
Schematic representation of all transketolase-catalysed reactions. Columns represent ketose-phosphates with m=5..8 carbons, while rows represent aldo-phosphates with n=3..6 carbons. Every possible reaction is indicated by an arrow, where the start and end squares stand for the substrates/products of the respective reaction.

**Figure 2B:**
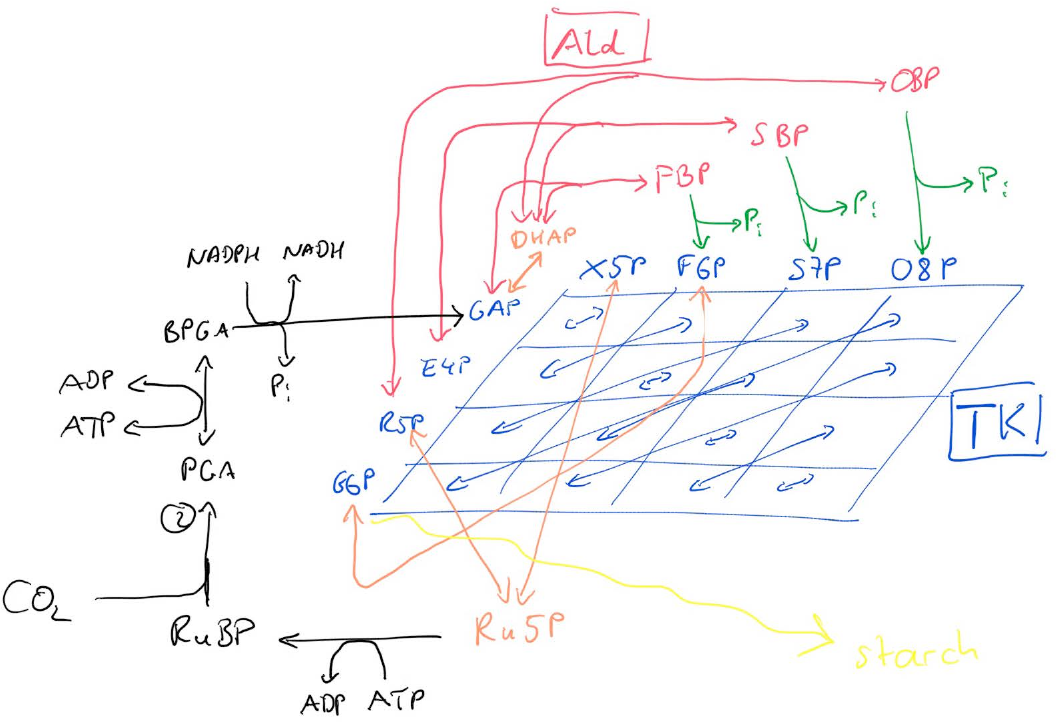
Alternative representation of the Calvin-Benson-Bassham Cycle. This representation includes the full complexity introduced by the broad substrate specificity of the transketolase and aldolase enzymes. In this representation the separation of the cycle into energy-driven (black and green) and entropy-driven (blue, red and orange) reactions becomes visible.

## Conclusions

What do the experimental and theoretical results discussed here tell us about the true structure of the CBB cycle? The preliminary results from mathematical models indicate that assuming the CBB cycle in its strict form is in accordance with the experimentally observed asymmetries of labels initially reported by Kandler and Gibbs. However, the results do certainly not preclude the existence of additional reactions, which possibly only have a minor influence on label distributions. Experimental evidence and rational arguments allow us to reinterpret the activity of the key enzyme transketolase. Instead of viewing it as catalysing exactly two reactions, we need to understand it as an enzyme catalysing a highly specific transfer of two carbons from a ketose phosphate to an aldose phosphate, but for a broad range of substrates. Since the reactions are highly reversible, the enzyme constantly mixes sugar phosphates of different lengths, much alike the ‘entropic’ carbohydrate-active enzymes experimentally and theoretically described by Kartal et al. in 2011 (49). The thermodynamic driving force is an increase in the mixing entropy of the substrate and product mixture. Because all of the 16 possible reactions are close to (or at) equilibrium, the individual forward and backward rates of the reversible processes must be considerably higher than the overall net conversion rate. Consequently, the scheme of the CBB cycle as originally proposed can still be considered as correct, understanding that it indicates the net conversion fluxes. However, it does ignore a large number of reversible transformations, which do not contribute to the net flux, but which certainly have some influence on the label incorporation dynamics. An attempt to depict the complexity caused by the reversible reactions with broad substrate specificity, catalysed particularly by transketolase (and to some degree by aldolase), is given in Fig 2B. Currently there is no possibility to determine the rates of the individual reactions separately *in vivo*. Fortunately, mathematical models are ideally suited to test how all these ‘invisible’ reactions influence predicted labelling patterns. Like the discovery of radioactive carbon isotopes provided scientists a new “eye” to look into processes inside leaves, so can mathematical models serve as yet another “eye”, by which internal details of metabolism can be made visible, that lie beyond our experimental observation capabilities.

